# A multiscale analysis of liver lobule fibrosis and its impact on drug propagation and metabolism – a DLA approach

**DOI:** 10.64898/2026.08.28.747748

**Authors:** Vahid Rezania, Dennis Coombe, Jack Tuszynski

## Abstract

Employing DLA methods, this paper explores the self-assembly of collagen fibers and resulting fibrosis at three scales up to the scale of regular lobule models. This allows a mechanistic exploration of the effects of collagen on drug transport (flow and diffusion) and metabolism. In addition, this method permits an analysis of fiber growth characteristics. First, variations of the DLA method of Parkinson et al (1994) will be used to generate multiple explicit collagen microfibril self-assembly using DLA particles in one dimension using cubic grid blocks of (4 mm)^3^ in a 240 × 20 × 20 grid model. The second stage will be to assess the consequences of various densities of these fibers in three dimensions on flow reductions at a higher scale. Here we utilize DLA methods in cubic grid blocks of (80 nm)^3^ to mimic 3D collagen self-assembly of fibrils. We then apply a pressure gradient or specified flow rates across a spatially gridded version of these models to quantify flow effects. This region represents a local zone of liver tissue affected by fibrosis. Analytic models of fibrotic effects on flow are employed for comparison. A third stage explores the implications of fibrosis in a liver lobule model using multiple grid blocks of size 3200 mm to represent the lobule tissue. Here, a continuum model of fiber density is employed, based on the previous two scales. The model also includes the effects of additional grid blocks representing sinusoidal flow paths found in the lobule. We contrast and quantify drug propagation and metabolism of molecular dissolved versus nanoparticle delivery vehicles in fibrotic media, achieved by upscaling explicit collagen distributions to appropriate average values.

## 1. Introduction

Previously, in Rezania et al (2014) we have developed flow simulation methods to quantify flow and metabolism in a liver lobule with DLA generated sinusoid paths traversing liver tissue using a heterogeneous model with grid blocks of size (3μm)^3^, Subsequently, Rezania et al (2024) have employed the same scale or larger grid size scales to estimate the effects of fibrosis, with assumed fibrotic flow reductions based on DLA fibrosis methods. This current paper proposes to use multi-scaling methods to better quantify these postulated flow reductions. Here, we first utilize two scales (4 nm and 80 nm, respectively), in a stagewise process to characterize this self-assembly. Table 1 summarizes the basic geometric parameters for each case.

**Table 1.** Geometrical and fluid tissue properties: two scales (micro vs mini scales)

| parameter | **20 × 20 × 240** | **40 × 40 × 40** |
| --- | --- | --- |
| $x,y,z$ (e-4 cm) | 4 nm | 80 nm |
| Gross formation volume (cm<sup>3</sup>) | 6.1440e-06 | 3.2768e-11 |
| Formation pore volume (cm<sup>3</sup>) | 1.4635e-06 | 7.8054e-12 |
| Aqueous phase volume (cm<sup>3</sup>) | 1.4635e-06 | 1.9513e-12 |
| Oil phase volume (cm<sup>3</sup>) | 0.0000e-06 | 0.5854e-12 |
| Tissue porosity: $f_{tis}$ | 2.3820e-01 | 2.3820e-01 |
| Tissue permeability (cm<sup>2</sup>): $K_{tis}$ | 7.3500e-10 | 7.3500e-10 |
| Flow rate (cm<sup>3</sup>/min) | 1.000e-05 | 7.000e-10 |

(1 Darcy = 0.9869e-12 m<sup>2</sup> = 0.9869e-8 cm<sup>2</sup> in engineering permeability units)

Collagen is a triple helix dominated by the amino acid glycine, proline, and hydroxyproline. Basic molecular tropocollagen forms a triple helix of amino acids of total length 300 nm and 1.5 nm diameter. Other useful properties include a molecular weight of 300 kDa and dehydrated density of 1.4 gm/cm^3^ (or specific volume of 0.73 cm^3^/gm).

Numerous studies of the self-assembly of type I collagen, rod-like proteins, to form elongated fibrils have been conducted. These fibrils vary in length from 300-1000 nm with diameters from 20-200 nm. Further aggregation of these fibrils into larger collagen fibers can then occur.

Earliest microfibril packing theories propose a staggered stacking of 5 units over integral multiples of a repeat distance of *d* = 67 nm, with a quasi-hexagonal packing perpendicular to this major axis (Hulmes and Miller, 1979 and Hulmes et al, 1995). Orgel et al (2006) presented a comprehensive analysis of microfibril formation using a cylinder model of tropocollagen plus a simple binding rule with a 3.4 *d* overlap coiling adjacent molecules into concentric layers. Silver et al (2003) investigated experimentally collagen self-assembly.

Buehler (2006 a, b) developed computational approaches to microfibril self-assembly based on advanced molecular dynamics models that allowed upscaling to continuum mechanical modelling. Vesentini et al (2013) summarize aspects of these calculations, including visualization of these self-assembly concepts (their figures 6 and 7). Our Fig 1 shows a similar schematic of this.

**Figure 1a.**
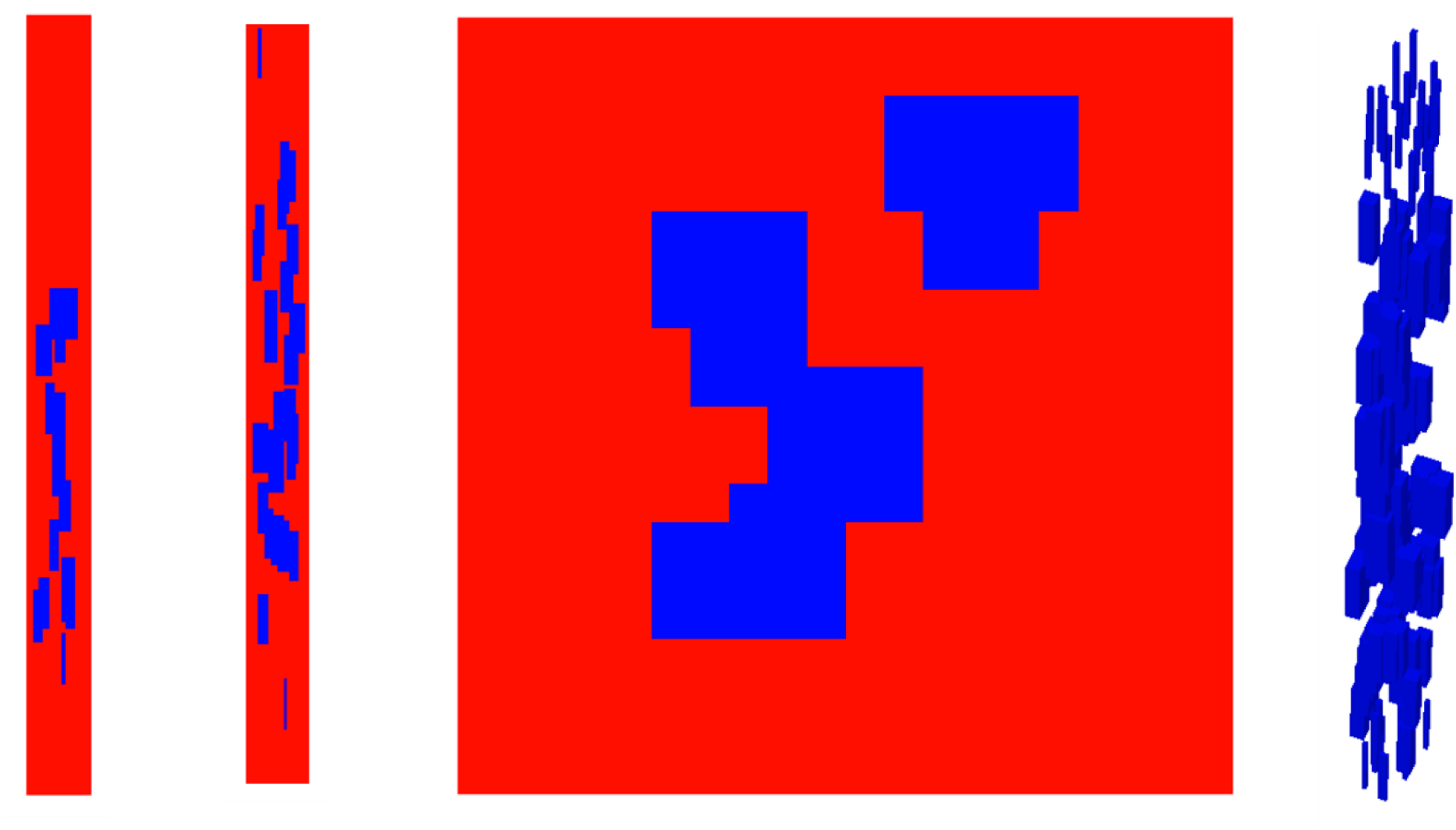
A 20 × 20 × 240 DLA microfibril formation from 0.781e+3 microfibrils with occupancy density of 0.123: red – normal tissue and blue – microfibrils. From left to right, *xz*-, *yz*-, and *xy*-planes at *y* = 10, *x* = 10, *y* = 120, and a 3D representation, respectively. In the 3D representation, the normal tissue sites have been removed for clarity.

**Figure 1b.**
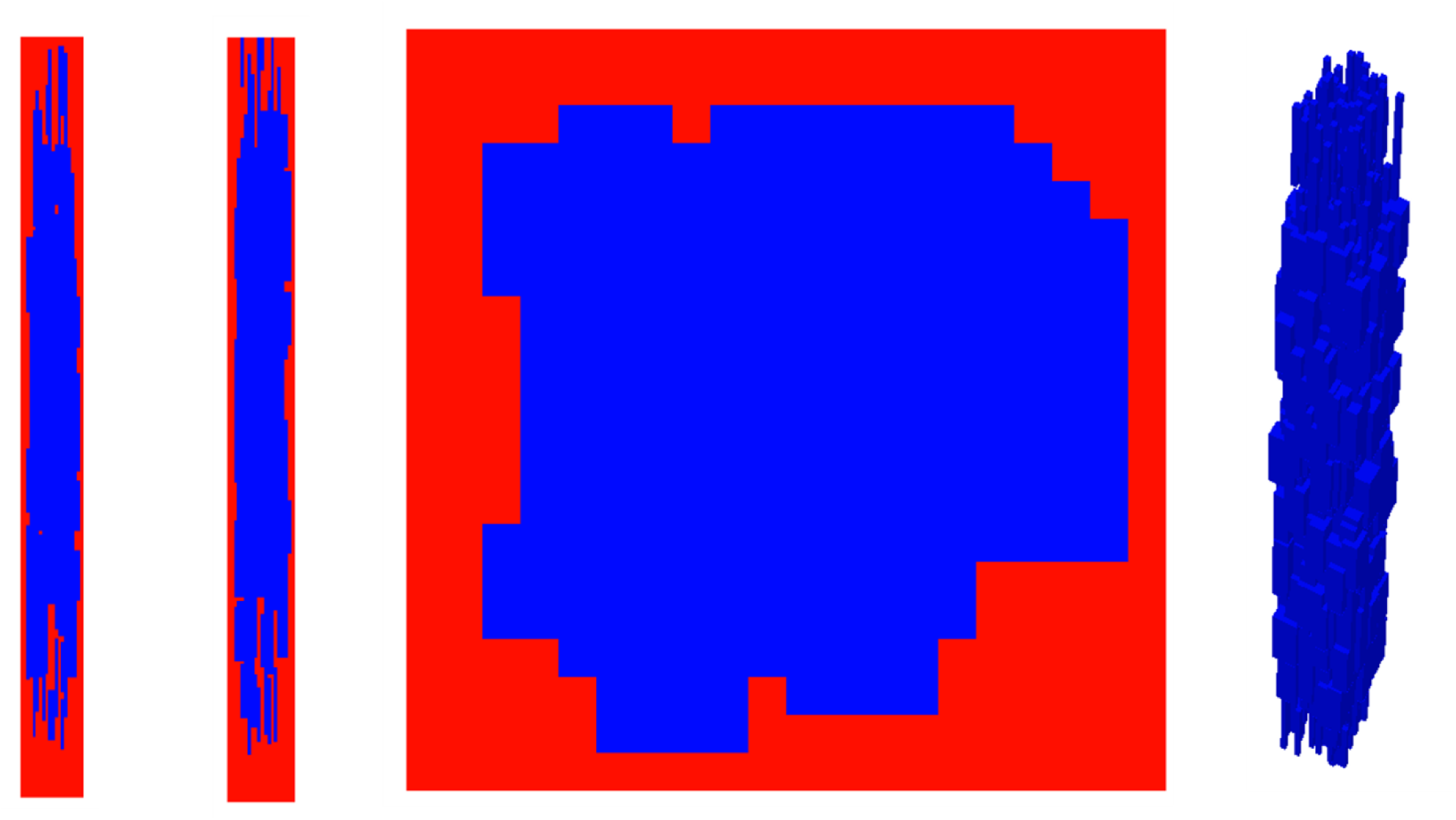
Same as Figure 1a but with 2.641e+3 microfibrils and occupancy density of 0.440.

We summarize the relevant properties by defining our terms:

a. tropocollagen - or collagen molecule (300 nm × 1.5 nm × 1.5 nm)
b. microfibril repeat unit used as DLA particle (64 nm × 4 nm × 4 nm)
c. fibril (960 mm × 80 nm × 80 nm), composed of DLA aggregated particles. Up to 10,000 particles were used to construct 100 or more microfibrils, depending on the desired fiber density.
d. collagen fibers of various sizes, (1D or 3D constructs of fibrils) found in multiple tissue types

This paper then investigates the dynamic consequences of such self-assembly on drug distributions.

## 2. Microfibril formation via collagen deposition- DLA approach

A model of collagen fibrogenesis based on DLA has been proposed by Parkinson et al (1994a, 1994b), later generalized to include stress responses by Parkinson et al (1997). These models again recognized the staggered stacking with repeat distance of *d* = 67 nm and generated structures of collagen fibrils including the elongated morphology and tip growth observed in experiments. Fractal dimensions and aggregate densities are estimated with this method. The size of the microfibrils generated are limited by the lattice grid size used (approximately 16 nm) and computer limitations. Note they use an 18 × 1 × 1 grid model of a collagen molecule and aggregate in one direction only. There are issues with their choice of grid representation of collagen, however. Note, as they mention, this molecule has dimension of 300 nm × 1.5 nm × 1.5 nm, so choosing a grid size of 1.5 nm, means their molecule should be 200 × 1 × 1, not 18 × 1 × 1.

We follow this DLA method here in part but recognize the necessary scale dependence of the desired fibrotic effects. Instead, we propose representing our collagen entity as the repeat unit of the collagen microfibril (involving parts of 5 collagen molecules associated together). The repeat unit length is 67 nm and a cross-section area of approximately 3nm × 3nm. This is a well-defined concept of repeat unit, see Figures 6 and 7 of the paper of Vesentini et al (2013). With our proposed grid (4nm), the repeat unit that we continually add in DLA is 22 × 1 × 1 grid point size. Note because a complete collagen molecule is 300 nm long, our starting point for DLA is to utilize 5 repeat units long i.e. 66 nm × 5 = 330 nm. Following the idea of Garcia-Ruiz and Otalora (1991), which was also employed by Parkinson et al, we explore an additional DLA step allowing aspects of “surface diffusion” of DLA particles to more stable growth sites after microfiber deposition.

Figures 1, S1 and S2 (in Supplementary Material) illustrate the resulting growth of 1D microfibrils, such that high densities almost completely fill the gridded area. We define the fibber occupancy density as the ratio of the total occupied sites to the total number of sites. With this grid model, we calculate the reduced effective porosity and 1D permeability with STARS by a flow calculation, see Table 2.

**Table 2.** Fibrotic effects on 1D flow (20 × 20 × 240 grid)

| **# Fibers** | **Fiber**<br>**occupancy**<br>**density** | **Volume** | **Volume**<br>**change**<br>**fraction** | **Flow** | **Flow**<br>**reduction** |
| --- | --- | --- | --- | --- | --- |
| 0.0 | 0.0 | 6.114e-06 | 0.0 | 1.00e-5 | 1.0 |
| 0.738e+3 | 0.1230 | 5.388e-06 | 0.8813 | 0.86e-5 | 0.86 |
| 2.070e+3 | 0.3449 | 4.025e-06 | 0.6583 | 0.54e-5 | 0.57 |
| 2.612e+3 | 0.4354 | 3.469e-06 | 0.5674 | 0.47e-5 | 0.47 |
| 3.474e+3 | 0.5791 | 2.586e-06 | 0.4230 | 0.35e-5 | 0.36 |
| 3.755e+3 | 0.6258 | 2.299e-06 | 0.3760 | 0.33e-5 | 0.33 |
| 3.874e+3 | 0.6457 | 2.177e-06 | 0.3561 | 0.32e-5 | 0.32 |
| 3.927e+3 | 0.6546 | 2.122e-06 | 0.3471 | 0.31e-5 | 0.31 |

Further refinements of our DLA aggregation method could be considered, based on later DLA method enhancements (Rothenbuhler et al, (2009); Nicolas-Carlock et al, (2016)). We expect, however, that the resultant basic flow effects will remain unchanged.

## 3. 3D Fibril formation via collagen deposition- DLA approach

Here, we downsize a single 3mm × 3mm × 3mm grid block from our original single lobule DLA model (Rezania et al, 2013) into a 40 × 40 × 40 grid model of 80 nm size blocks. This would allow DLA to model individual collagen-like particles following the 3D model of Parkinson et al (1994) but developed at a larger scale than theirs.

Our implicit 3mm × 3mm × 3mm grid block consists of distributions of 3D DLA generated fibrils (e.g., *N* fibrils generated perpendicular to the *x*-direction (of 80 nm diameter) plus *N* fibrils in the *y*-direction and *N* fibrils in the *z*-direction) in our 40 × 40 × 40 grid cell model. Figures 2 and S3‒S5 (see Supplementary Material) illustrate the resulting growth of 3D microfibrils, such that high densities almost completely fill the gridded area. With this grid model, we calculate the reduced effective porosity and 3D permeability with STARS by a flow calculation, see Table 3.

**Figure 2a.**
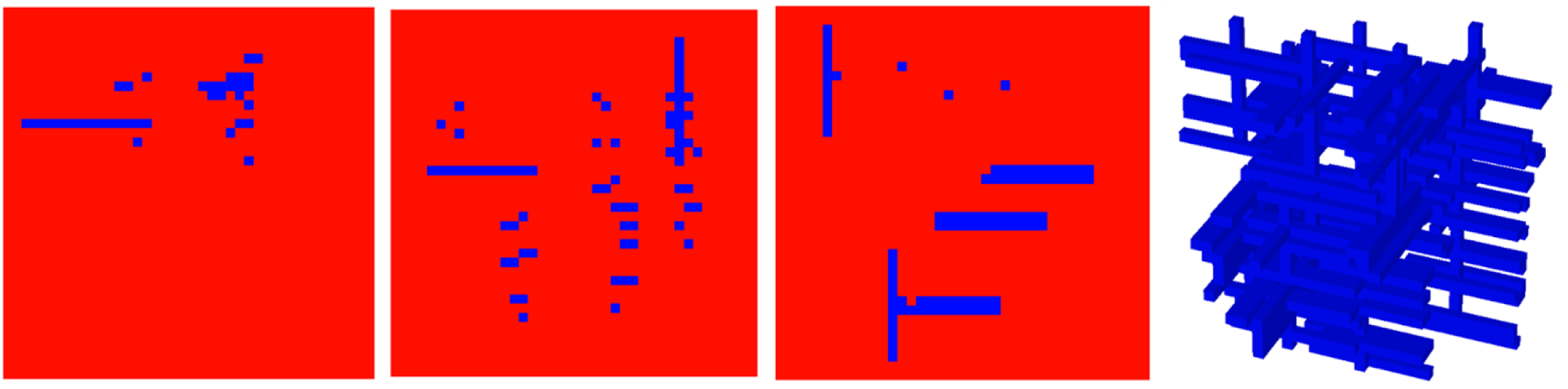
A 40 × 40 × 40 DLA fibril network: fibril occupancy density 0.031 (164 fibrils): red – normal tissue and blue – fibril network. From left to right, *xz*-, *yz*-, and *xy*- planes at *y* = 20, *x* = 20, *z* = 20, and a 3D representation, respectively. In the 3D representation, the normal tissue sites have been removed for clarity.

**Figure 2b.**
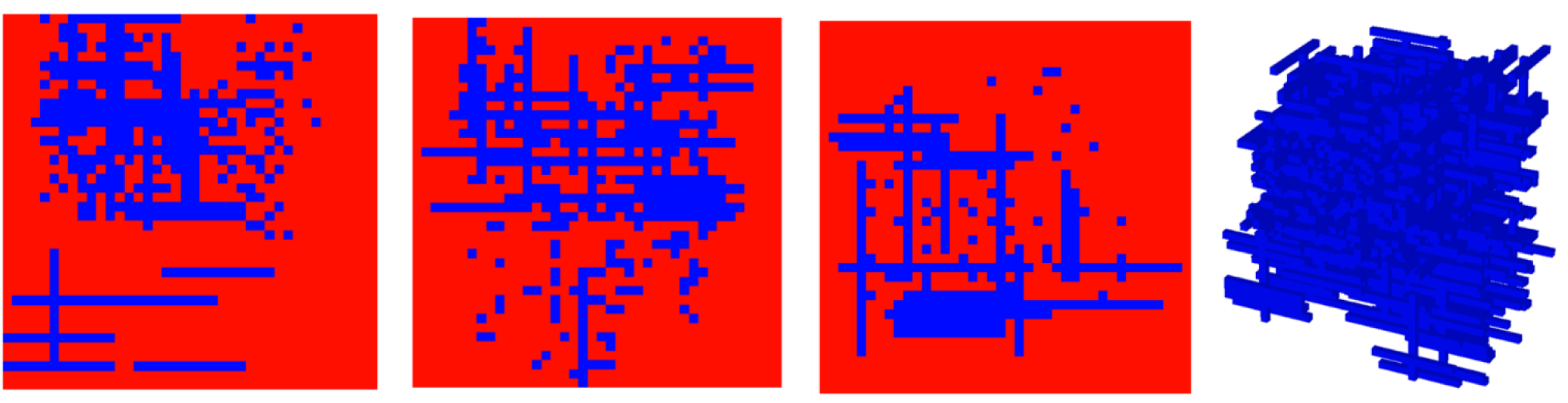
Same as Figure 2a with fibril density 0.189 (1007 fibrils).

**Table 3.** Fibrotic effects on 3D flow (40 × 40 × 40 grid)

| **# Fibers** | **Fiber**<br>**occupancy**<br>**density** | **Volume** | **Volume**<br>**change**<br>**fraction** | **Flow** | **Flow**<br>**reduction** |
| --- | --- | --- | --- | --- | --- |
| 0.0 | 0.0 | 3.277e-11 | 0.0 | 70e-11 | 1.0 |
| 1.065e+3 | 0.1996 | 2.623e-11 | 0.8000 | 39e-11 | 0.557 |
| 2.589e+3 | 0.4854 | 1.686e-11 | 0.5144 | 19e-11 | 0.271 |
| 3.123e+3 | 0.5854 | 1.358e-11 | 0.4144 | 15e-11 | 0.214 |
| 4.103e+3 | 0.7692 | 0.7562e-11 | 0.2307 | 9.0e-11 | 0.129 |
| 4.331e+3 | 0.8120 | 0.6159e-11 | 0.1879 | 8.4e-11 | 0.120 |

As a simple reference comparison with our approach, Pederson et al (2007) have postulated a regular collagen deposition pattern, assuming 700 nm diameter collagen microfibrils with a 4 mm spacing between fibrils in 3D on a 32 mm × 16 mm × 16 mm model. This resulted in a porosity reduction of *φ* = 0.934 and a calculated permeability of approximately 2e-8 cm^2^ = 2000 mD. Stein et al (2007) and Laing et al (2013) have presented experimental characterization of random fiber distributions.

## 4. Fibrotic effects on permeability

Theories on flow in fibrous porous media have been discussed by many authors, basically correlating permeability change with porosity change. Conceptually, this porosity change with fibrosis is related to extra collagen deposition.

Theories on the permeability of fibrous materials and comparison with experiments have been reviewed by Jackson and James (1986) and Zhu et al (2017). Most theories start with the premise of regular fiber alignment and consider flows parallel or perpendicular to these fibers. As emphasized by Jackson and James, an isotropic 3D fiber model can be developed by combining these parallel and perpendicular results using a 1/3 and 2/3 ratio, as for example

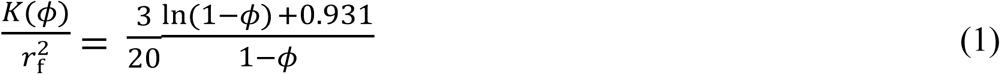

Standard porous media flow correlations relating porosity and permeability include the Kozeny-Carmen equation (Bear, (1972), as used in STARS) originally developed to model packed beads and sand grains

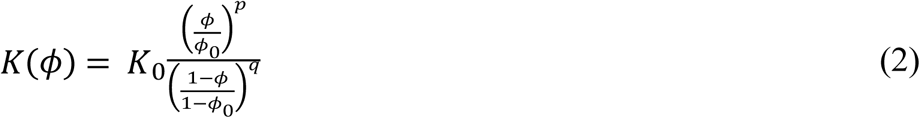

Here, *K*_0_ represents the square of hydraulic radius and *p* and *q* are adjustable powers. Typically, *p* = 3 and *q* = 2. Interestingly, Costa (2006) and Zhu et al (2017) both emphasize that CK-type correlations can usefully be applied to fibrous beds as well, with Costa examining fractal effects as well. We will assume this consistency and apply CK models in our calculations, as all models presented express permeability as a function of porosity (collagen volume fraction) and measures of fiber radius.

Table 4 summarizes the porosity/permeability results of the DLA fibril effects (see Table 3), idealized to three characteristic regions of the lobule (sinusoid fibrosis, implicit tissue fibrosis, explicit tissue fibrosis) as proposed in our earlier lobule model with fibrosis (Rezania et al, 2024). These permeability values will be employed for our quantitative simulations on drug propagation in section 6. Interestingly, Costa’s generalized CK equation (his equation 17) seems to approximately map our calculated DLA permeability reductions using “*p* = 1 + *n*” and “*q* = *n*” with low “*n* < 1” values, reflecting significant fractal effects on specific surface areas.

**Table 4.**
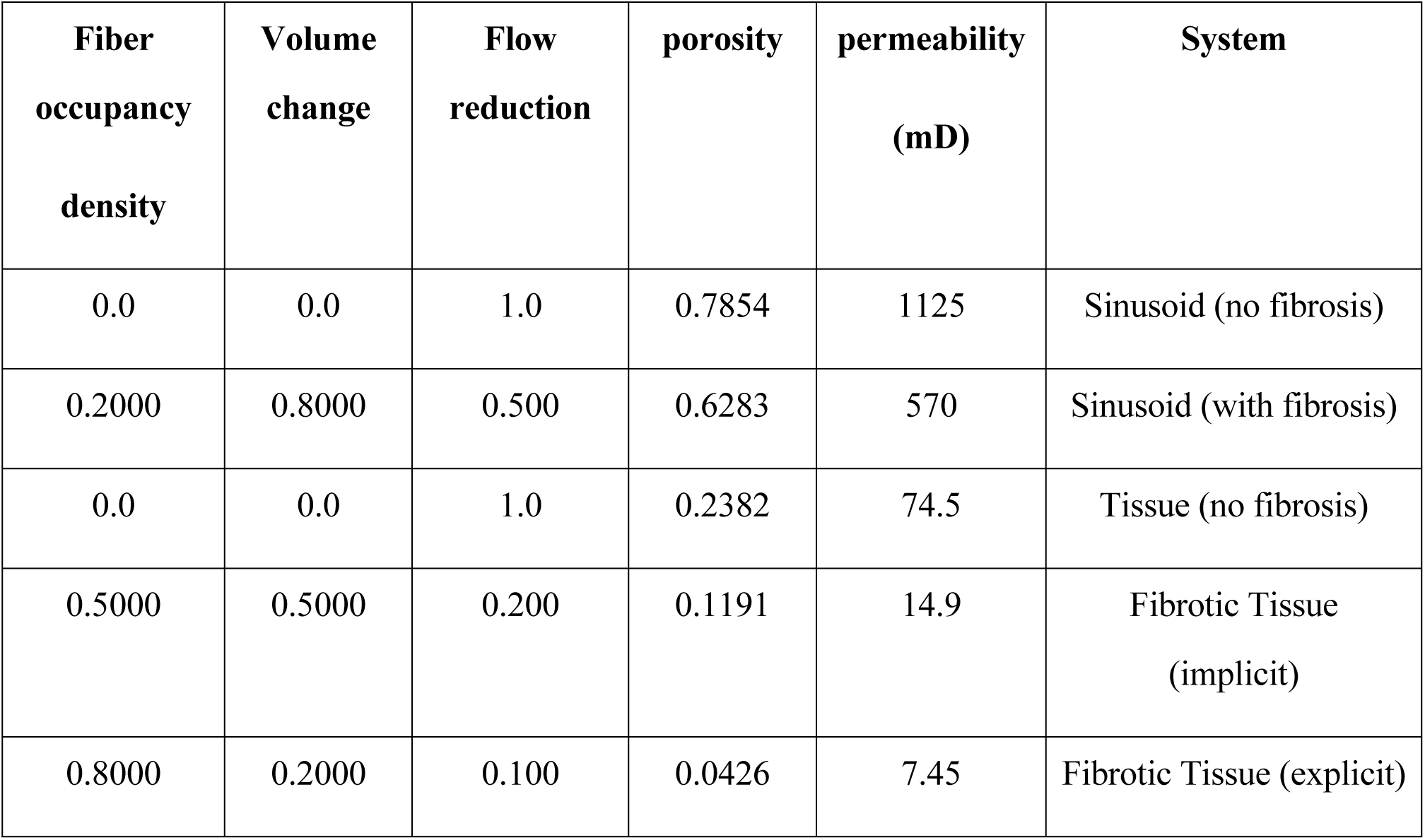
Fibrotic effects on flow (upscaled grid)

| **Fiber**<br>**occupancy**<br>**density** | **Volume**<br>**change** | **Flow**<br>**reduction** | **porosity** | **permeability**<br>**(mD)** | **System** |
| --- | --- | --- | --- | --- | --- |
| 0.0 | 0.0 | 1.0 | 0.7854 | 1125 | Sinusoid (no fibrosis) |
| 0.2000 | 0.8000 | 0.500 | 0.6283 | 570 | Sinusoid (with fibrosis) |
| 0.0 | 0.0 | 1.0 | 0.2382 | 74.5 | Tissue (no fibrosis) |
| 0.5000 | 0.5000 | 0.200 | 0.1191 | 14.9 | Fibrotic Tissue<br>(implicit) |
| 0.8000 | 0.2000 | 0.100 | 0.0426 | 7.45 | Fibrotic Tissue (explicit) |

## 5. Fibrotic effects on diffusion

Reactive drug transport requires the additional consideration of diffusive flows and reactions. These effects are typically characterized by two dimensionless numbers: the Peclet number describing the ratio of convective to diffusive transport, and the Damkohler number describing the ratio of reactive rates to diffusive transport (Bear, 1972). Resultant observed behavior can vary widely when these dimensionless variables change significantly. Changing fibrosis effects can impact these processes at multiple scales as we discuss next.

Effective diffusion is impacted by the presence of tissue fibers and depends on three factors – concentration of fibers *c*_f_ = (1 ̶ *ϕ*), fiber radius *r*_f_ (often estimated as square root of tissue permeability), and molecule size *r*_s_ (molecular radius). Ogston et al (1973) gave a first representation of such effects with *l* = *r*_f_/*r*_s_

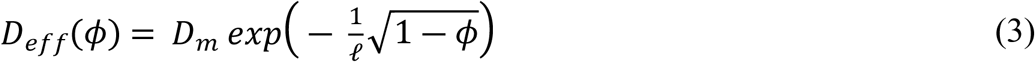

where *D*_m_ is molecular diffusion constant in water or blood. Here fiber size *r*_f_ is usually expressed as (*K*/*ϕ*)^0.5^ where the permeability is also a function of porosity as we discussed previously.

Equation (3) is an example of stretched exponentials, accounting for steric (or tortuosity) effects. More complex realistic models also account for hydrodynamic contributions as a separate factor (Johnson et al, 1996; Clague and Phillips, 1996). First estimates of this factor used effective medium theory, resulting in a power law expression with *r*_s_ /*Κ* ^0.5^. An alternate approach (Clague and Phillips, 1996) correlated this factor with a separate stretched exponential term

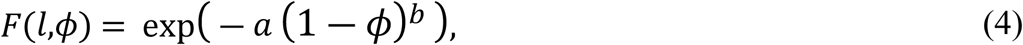

with both factors (*a* and *b*) expressed as power series in l = *r*_f_/*r*_s_, (Amsden, 1998; Phillips, 2000). The resulting complete expression for hindered diffusion involves two stretched exponential factors and typically provides useful matches with experiment

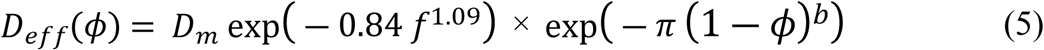

Here,

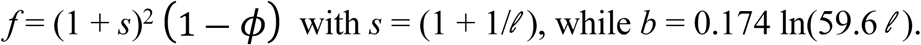

All models presented express diffusion as a function of porosity (collagen volume fraction) and ratios of molecular radius / fiber radius (here called, *s*) and/or fiber radius / molecular radius (here called, *l*). Finally, using ratios of these expressions for different fiber volume fractions allows estimations of changing levels of fibrosis on diffusion.

Stylianopoulos et al (2010) theoretically explore the diffusion of macromolecules and nanoparticles through stochastically-generated fiber networks of fixed fiber size (100 nm) but various concentrations and orientations. They utilize a random walk algorithm and compare their results to the Amsden expression, equation (5).

Table 5 illustrates the analytic porosity/diffusion results of fibrosis based on the Amsden equation (our equation 5) for the basic case of Au paclitaxel NP (Gao et al, 2013) of size 80 nm diffusing through a microfiber network of 80 nm. Table 6 then utilizes these porosity/diffusion results idealized to three characteristic regions of the lobule (sinusoid fibrosis, implicit tissue fibrosis, explicit tissue fibrosis) as proposed in our earlier lobule model with fibrosis (Rezania et al, 2024). These diffusion values will be employed for our quantitative simulations on drug propagation in section 6 below.

**Table 5.** Fibrotic effects on diffusion constant (analytic Gao et al NP)

| **Fiber**<br>**occupancy**<br>**density** | **Porosity**<br>**change** | **$D/D_0$** |
| --- | --- | --- |
| 0.0 | 1.0 | 1.0 |
| 0.1 | 0.9 | 0.3984 |
| 0.2 | 0.8 | 0.1904 |
| 0.3 | 0.7 | 0.0945 |
| 0.4 | 0.6 | 0.0479 |
| 0.5 | 0.5 | 0.0246 |
| 0.6 | 0.4 | 0.0127 |
| 0.7 | 0.3 | 0.0066 |
| 0.8 | 0.2 | 0.0035 |
| 0.9 | 0.1 | 0.0018 |

Sinusoid diffusion constant (no fibrosis), $D_{0s} = 4.22\text{e-}6 \text{ mm}^2/\text{sec} = 2.53\text{e-}6 \text{ cm}^2/\text{min}$ (Gao et al, 2013)
Tissue diffusion constant (no fibrosis), $D_{0t} = 1.89\text{e-}6 \text{ mm}^2/\text{sec} = 1.13\text{e-}6 \text{ cm}^2/\text{min}$ (Gao et al, 2013)

**Table 6.** Fibrotic effects on Gao et al’s diffusion constant (upscaled grid)

| **Fiber occupancy**<br>**density** | **Porosity**<br>**change** | **$D/D_0$** | **$D$**<br>**(cm<sup>2</sup>/min)** | **System** |
| --- | --- | --- | --- | --- |
| 0.0 | 0.0 | 1.0 | 2.53e-6 | Sinusoid (no<br>fibrosis) |
| 0.2000 | 0.8000 | 0.1904 | 4.82e-7 | Sinusoid (with<br>fibrosis) |
| 0.0 | 0.0 | 1.0 | 1.13e-6 | Tissue (no<br>fibrosis) |
| 0.5000 | 0.5000 | 0.0246 | 2.78e-8 | Fibrotic Tissue<br>(implicit) |
| 0.8000 | 0.2000 | 0.0035 | 3.96e-9 | Fibrotic Tissue<br>(explicit) |

Sinusoid diffusion constant (no fibrosis), $D_{0s} = 4.22\text{e-}6 \text{ mm}^2/\text{sec} = 2.53\text{e-}6 \text{ cm}^2/\text{min}$ (Gao et al, 2013)
Tissue diffusion constant (no fibrosis), $D_{0t} = 1.89\text{e-}6 \text{ mm}^2/\text{sec} = 1.13\text{e-}6 \text{ cm}^2/\text{min}$ (Gao et al, 2013)

## 6 Fibrotic effects on drug propagation

Here, we investigate the role of fibrosis on the propagation and reactivity of drug molecules in a liver lobule using our upscaling methodology. More specifically we will compare the distribution and reactivity of (essentially) molecularly dissolved paclitaxel to that of a paclitaxel-containing nanoparticle of size 80 nm, in a collagen-enhanced tissue also of fiber size 80 nm. Here, we use the common drug formulation Taxol to represent “molecularly dissolved” form of paclitaxel.

Our computational lobule model is based on earlier work (Rezania et al, 2013; Rezania et al, 2024) utilizing DLA techniques to generate fibrosis patterns at the lobule scale. As such, this represents a third (continuum) scale of fibrosis utilizing DLA techniques, and the resulting distributions are consistent with that seen experimentally (Gole et al, 2023). Here we differentiate between implicit fibrosis levels (fiber sizes smaller than our selected grid sizes) and explicit fibrosis (fiber sizes extending overt this size). Figure 2 of Gole illustrates this size distribution experimentally. Furthermore, these models employ sinusoid patterns also generated via a DLA technique. All DLA factors affect flow, diffusion, and metabolism, as we shall show, and vary layer by layer. It is emphasized that all models are fully 3D versions of a lobule, although we often extract 2D areal plots to illustrate spatial distributions. To visualize distributions in 3D, we have chosen to employ grouped layer cutout representations. Table 7 summarizes the basic lobule characteristics of these models.

**Table 7.** Implicit and explicit fluid properties of a single DLA lobule (original vs fibrotic)

| Parameter | Original<br>values<br>(no fibrosis) | Fibrotic<br>(implicit) | Fibrotic<br>(explicit) |
| --- | --- | --- | --- |
| Tissue permeability (cm<sup>2</sup>): $K_{tis}$ | 7.3500e-10 | 7.3500e-11 | 7.3500e-12 |
| Tissue porosity: $f_{tis}$ | 2.3820e-01 | 2.3820e-01 | 2.3820e-01 |
| PAC/PAC-OH Diffusion (cm<sup>2</sup>/min): $D_0$ | 2.5000e-04 | 1.9162e-04 | 1.9162e-06 |
| PAC/PAC-OH Diffusion (cm<sup>2</sup>/min): $D_{tis}$ | 2.5000e-05 | 1.4945e-05 | 1.4945e-07 |
| O<sub>2</sub> Diffusion (cm<sup>2</sup>/min): $D_{O2-0}$ | 1.8000e-03 | - | - |
| O<sub>2</sub> Diffusion (cm<sup>2</sup>/min): $D_{O2-tis}$ | 1.2000e-03 | - | - |

(1 Darcy = 0.9869e-12 m<sup>2</sup> = 0.9869e-8 cm<sup>2</sup> in engineering permeability units)

Using the fibrosis levels proposed above, Figure 3 compares the permeability distributions, without and with fibrosis, utilizing a group layer cutout visualization of our 3D lobule model. Figure 4 illustrates the effects of fibrosis on the production behavior of a reactive small molecule O_2_, essentially indicating fibrosis delays O_2_ propagation.

**Figure 3:**
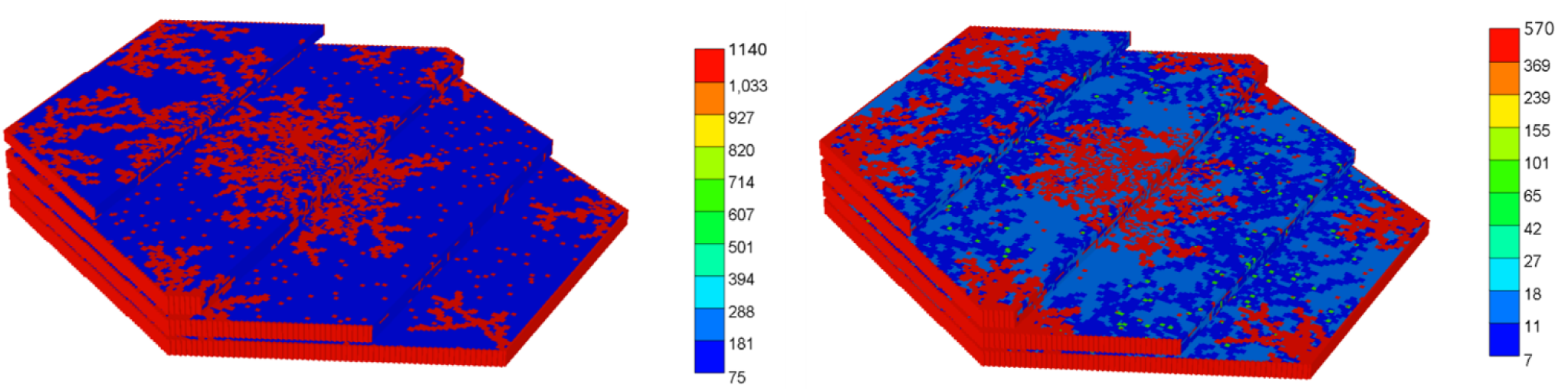
Lobule permeability distribution: (a) Left: no fibrosis (b) Right: with fibrosis.

**Figure 4:**
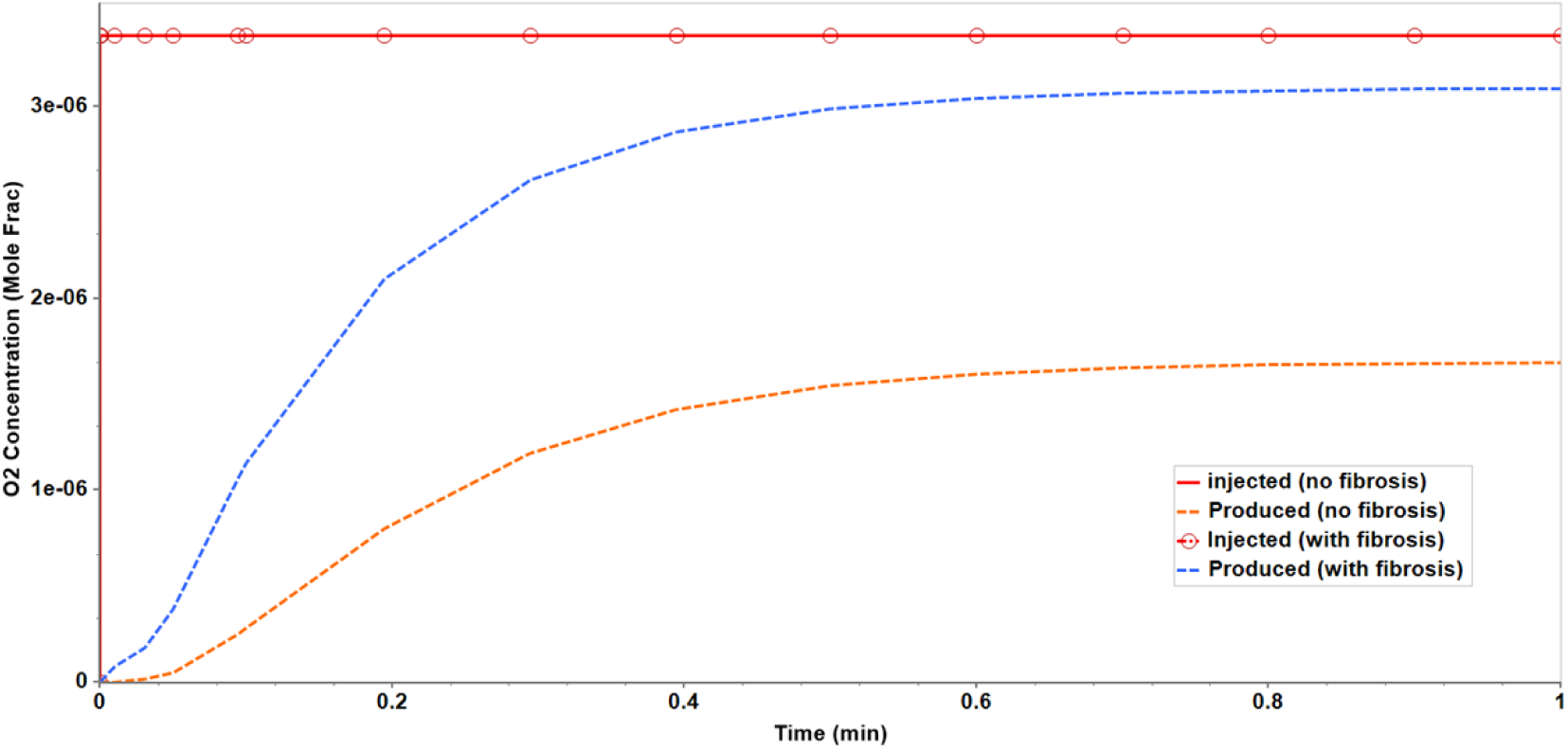
O_2_ production profile versus time: no fibrosis vs fibrosis.

Figure 5a compares the steady state O_2_ concentration distribution in these two cases for one typical layer, illustrating a substantial shift of O_2_ levels (higher with fibrosis), even though the O_2_ reaction kinetics parameters have not changed. This change in O_2_ distributions is extremely significant, as O_2_ distributions have been found to determine zonation behavior in the liver lobule (Keitzmann, 2017).

**Figure 5a.**
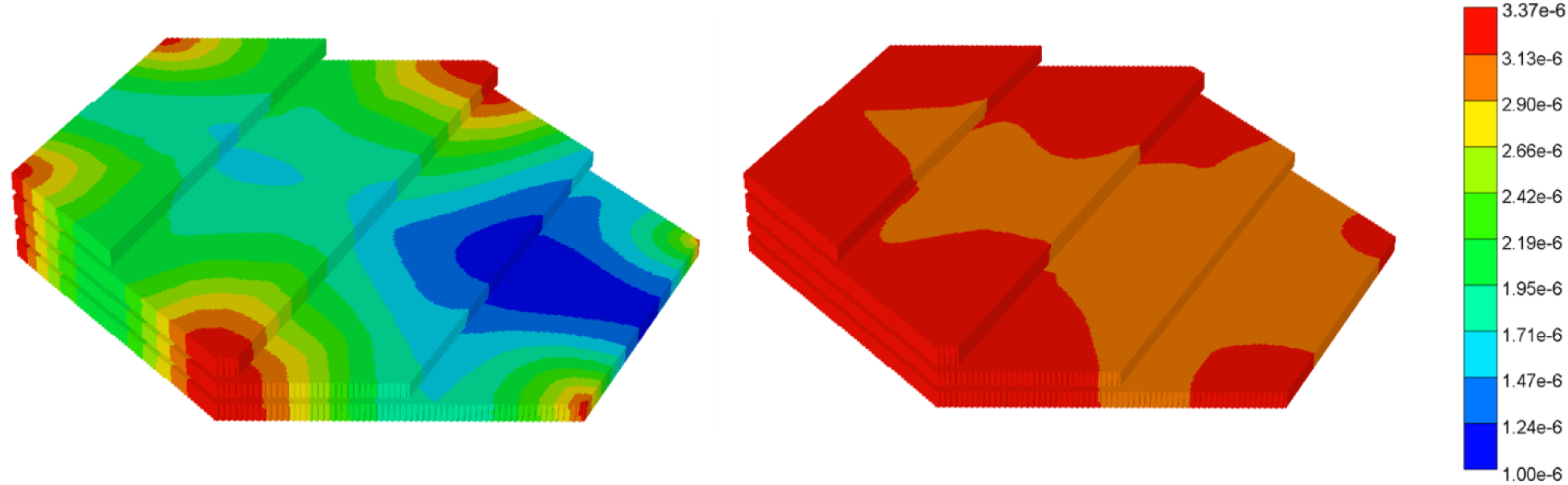
O_2_ concentration profile (2D slice with base scaling): (a) Left: no fibrosis, (b) Right: with fibrosis.

**Figure 5b.**
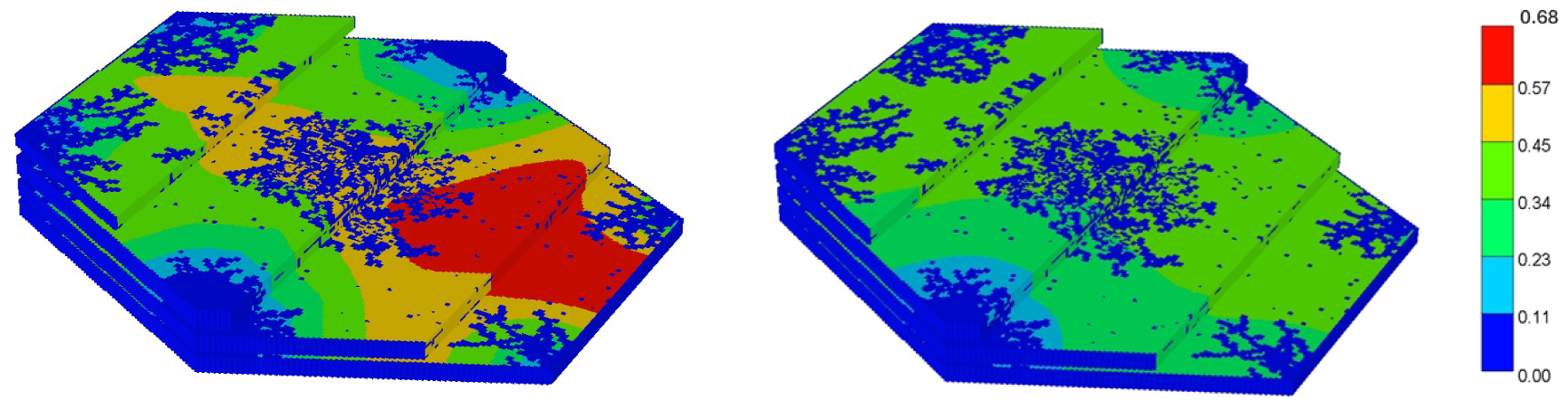
Resultant dimensionless CYP distribution (3D cutout plot with base scaling): (a) Left: no fibrosis, (b) Right: with fibrosis.

**Figure 6:**
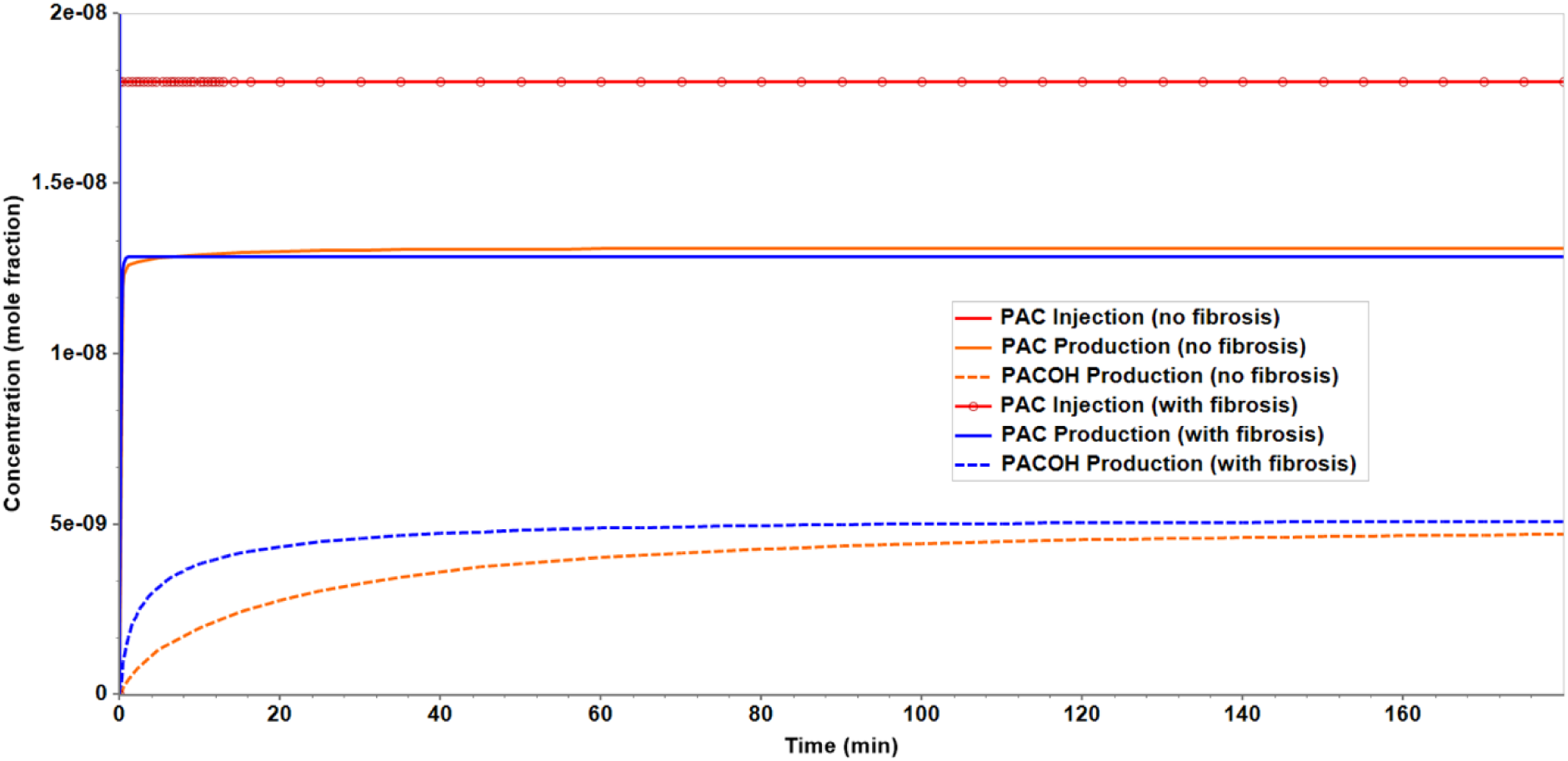
molecularly dispersed PAC and PACOH production profile versus time; without and with fibrosis.

**Figure 7:**
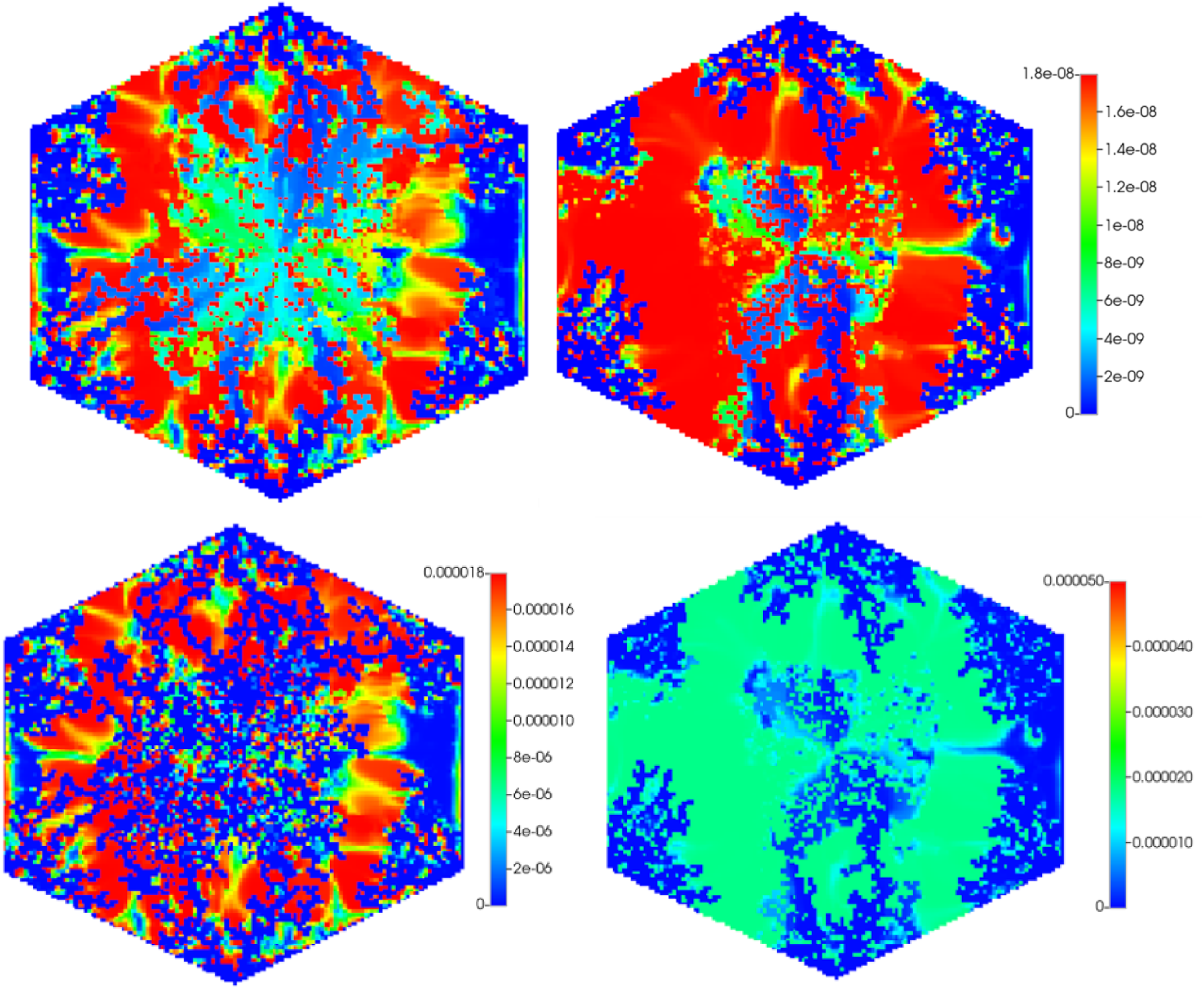
Spatial distribution of PACOH species in water (top panels) and on oil (bottom panels): Left: no fibrosis. Right: with fibrosis (LKV 1000)

The effect of fibrosis on zonation has been demonstrated experimentally (Ghallab et al, 2019). For our drug simulations given below, we will focus on the shift of metabolizing (CYP) enzyme distribution. Following our previous methods connecting O_2_ distributions (Figure 5a) with CYP distributions (Rezania et al, 2013, Rezania et al, 2024), the resulting CYP distributions (Figure 5b) are consistent with that found experimentally (Ghallab et al, 2019) – namely a lower region of pericentral enzymes with fibrosis.

Adjustment of the color scales and comparison of 2D, 3D representations allow further views of this process (see Figures S6‒S8 in Supplementary Material). Using the Rezania (2013 and 2024) models as a basis, we extend these in two ways to realistically compare paclitaxel drug propagation and reaction over a 3hour injection timeframe. First molecularly dissolved (i.e. Taxol-like) drug uptake over this time must account for paclitaxel protein binding both extracellularly and intracellularly. This has been experimentally analyzed by Au and coworkers (e.g. Fig. 1 and Table 1 from Kuh et al, 2000) and essentially results in a paclitaxel cell-to-medium distribution ratio of approximately 1000. This binding effect essentially slows paclitaxel propagation into tissue by the same factor. Free drug effects are thereafter modelled by our standard metabolic rate process, converting PAC to PACOH, based on Vaclavikova et al (2004) experiments. This process is assumed to be pericentrally-based in the absence of fibrosis, as seen experimentally.

Secondly, paclitaxel nanoparticle uptake kinetics are based on the experiments and model of Au and coworkers (Gao et al, 2013) for HSPC liposomes. This uptake model is analogous to that developed for influenza virus particles of similar sizes by Nir and coworkers (Nunes-Correia et al 1999). Here 4 required model parameters include rates of cell-NP association, cell-NP dissociation, and NP internalization, as well as an uptake saturation parameter describing the number of binding sites per cell. We furthermore assume the NP internalization leads directly to NP dissociation (PACNP to PAC), although this may involve a further rate delay. Once molecularly dissolved, PAC metabolism is governed by the previously described metabolic rate model, Vaclavikova et al (2004). Table 8 presents the resulting drug rate parameters employed for both drug cases.

**Table 8.** Drug partitioning and kinetic parameters for molecularly dissolved (Taxol-like) and nanoparticle paclitaxel cases.

| Parameter | Literature value | STARS units |
| --- | --- | --- |
| PAC, PACOH cell/aqueous partition coefficient | 1000<sup>a</sup> | 1000 |
| **PAC → PACOH metabolism:** |  |  |
| $V_m/K_m$ | 0.36/hr <sup>b</sup> | 6e-3 min<sup>-1</sup> |
| $K_m$ | 10 uM <sup>b</sup> | 1.8e-7 molefr |
| **NP kinetic uptake<sup>c</sup>:** |  |  |
| Cell_NP association rate | 5.5e+4 M<sup>-1</sup> s<sup>-1</sup> <sup>c</sup> | 1.8e+8 molfr<sup>-1</sup> min<sup>-1</sup> |
| Cell_NP dissociation rate | 3.4e-4 s<sup>-1</sup> <sup>c</sup> | 2.04e-2 min<sup>-1</sup> |
| NP internalization rate | 0.4e-4 s<sup>-1</sup> <sup>c</sup> | 2.4e-3 min<sup>-1</sup> |
| Cell binding maximum | 0.29e+6/cell <sup>c</sup> | 5.22e-11 molfr |

<sup>a</sup>: Kuh et al. (2000)
<sup>b</sup>: Vaclavikova et al. (2004)
<sup>c</sup>: Gao et al. (2013)

With these parameters in our given basic lobule model, four simulations for 3 hours of paclitaxel injection are conducted and analyzed:

i. molecularly dispersed (Taxol-like) paclitaxel in nonfibrotic and fibrotic lobule
ii. nanoparticle paclitaxel in nonfibrotic and fibrotic lobule.

Figures 6 summarizes the production characteristics of molecularly dispersed PAC injection, comparing without and with fibrosis cases. Here, it is seen that a little less than 1/3 of injected PAC is converted to PACOH over the 3-hour injection period. Fibrosis delays somewhat the process but otherwise has little effect on the produced PACOH levels. This contrasts to observations of drug propagation/reaction in spheroid experiments (Kuh et al, 2000) where little drug penetration is observed with fibrosis. In a lobule the presence of convective sinusoid flow paths still allows significant amount of lobule drug penetration and metabolism. It should be emphasized because of the assumed protein binding levels used in the model (LKV =1000), the produced PAC and PACOH levels observed in the aqueous phase are 1000 times lower than those internal cell levels.

Figure 7 compares the steady state spatial distribution of PACOH for a typical lobule layer, without and with fibrosis respectively. Both aqueous and cell PACOH concentrations are shown, which differ by the steady-state partitioning factor of 1000. The no-fibrosis plots with particular clearity show the division of zonation into zone1, zone2, zone3 areas of zonation. Fibrosis is seen to allow more pericentral penetration of PACOH, relative to the no-fibrosis case. These distributions reflect the changes in CYP enzyme zonation distributions caused by fibrosis (see Figure 5).

These results are contrasted to the nanoparticle injection of PAC described next. Figure 8 summarizes the production characteristics PACNP injection, comparing without and with fibrosis cases. Here it is seen that a minimal amount of injected PACNP is internalized by the liver cells over the 3-hour injection period. We emphasize that no PACOH product is produced by this time point, although some cell PACOH conversion occurs (see below). This NP distribution is behaving as intended, as the nanoparticle injection is designed for long circulation times in the human body.

**Figure 8a.**
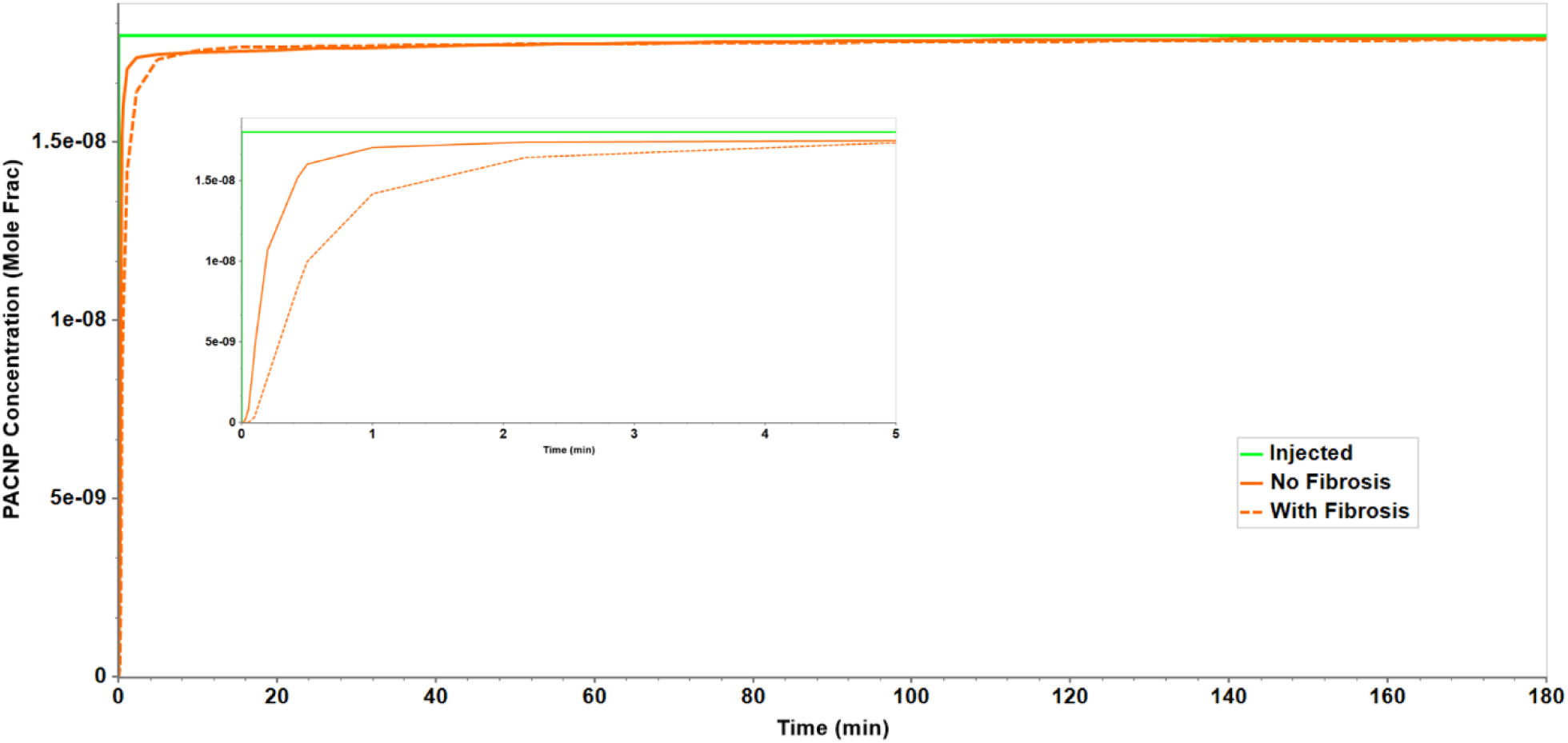
PACNP production profile versus time; no fibrosis and with fibrosis. The inset shows a magnified view of the first five minutes of the results.

**Figure 8b.**
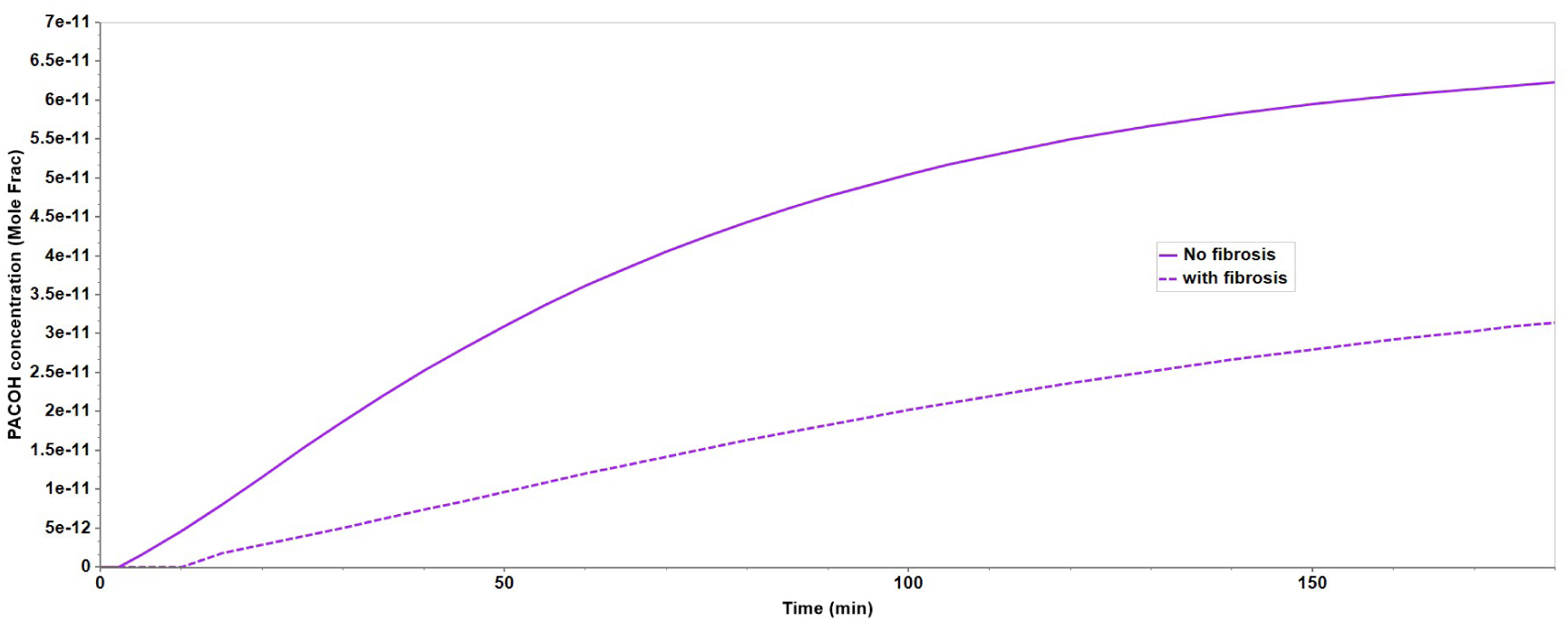
PACOH production profile versus time; no fibrosis and with fibrosis.

Fibrosis delays minimally the process of nanoparticle propagation through the lobule (see especially the insert profile of Figure 8a). The difference can be attributed to reduced nanoparticle diffusion with fibrosis. But as the process is still convection-dominated, this results in a small effect.

Figure 9 compares the spatial distribution of PACOH for a typical lobule layer, without and with fibrosis respectively. Both aqueous and cell PACOH concentrations are shown. While the no fibrosis aqueous concentrations reach injected concentrations in limited regions of the lobule, the predicted maximum PACOH concentration in the fibrotic case reaches approximately half this value. The cell concentrations of PACOH are here not yet at steady state values as shown and differ substantially between the no fibrosis and fibrotic cases. These distributions, while again reflecting the changes in CYP enzyme zonation distributions caused by fibrosis, show behavior dominated by kinetic aspects of metabolism. Nanoparticle uptake slows all aspects of the overall process.

**Figure 9:**
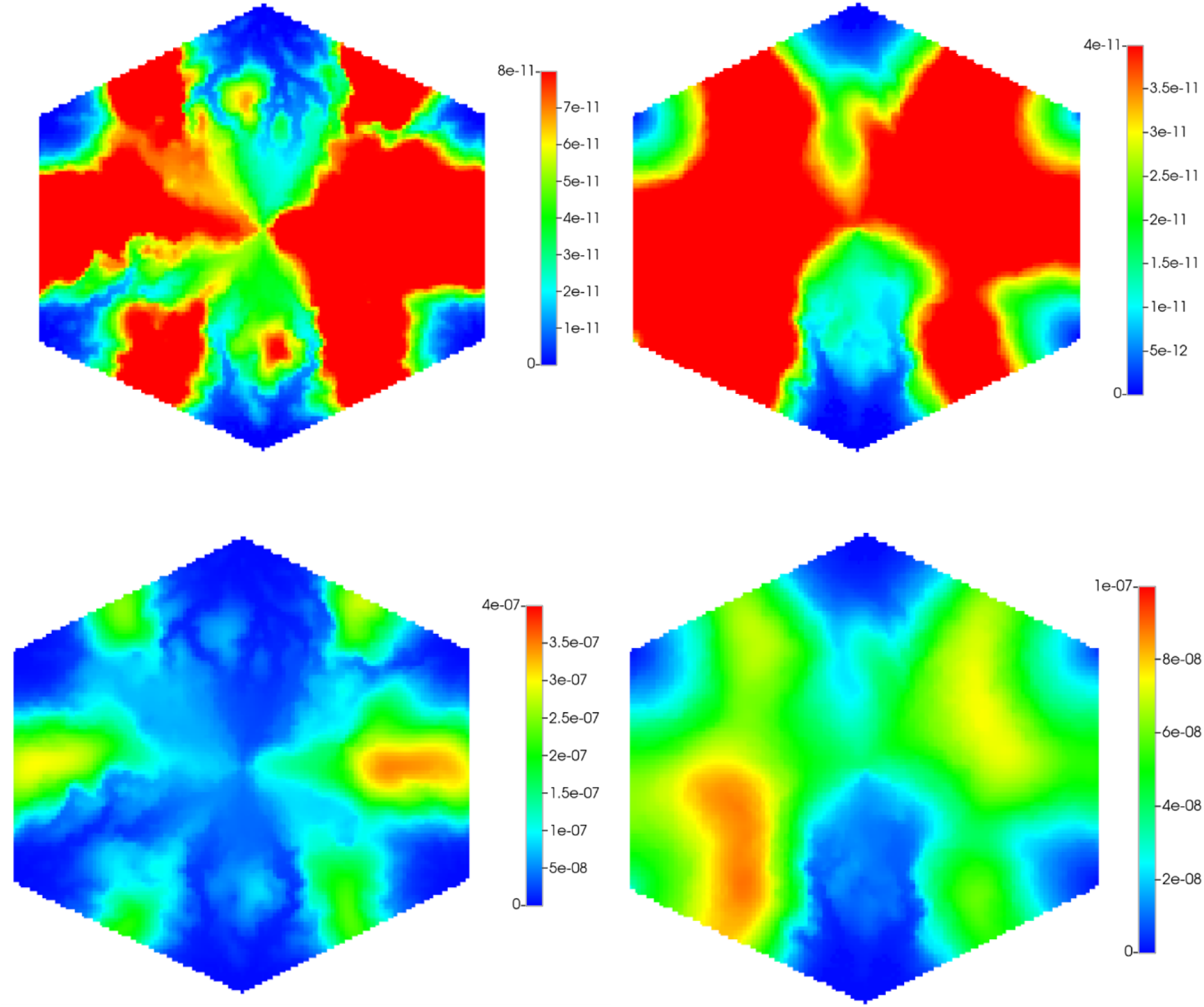
Spatial distribution of PACOH species in water (top panels) and on oil (bottom panels): Left: no fibrosis. Right: with fibrosis (NP)

Note while the PAC to PACOH metabolic enzymes are assumed to be zoned, our simulations assume no zonation for the nanoparticle uptake receptors (due to lack of experimental information). If these proteins also follow a peri-central dominated zonation, the observed zonation effect on PACOH would be amplified further.

## 7. Conclusion

This paper has demonstrated how to employ DLA scaling methods to estimate the effects of collagen fibrosis on liver tissue function at several scales. These estimates are shown to be consistent with imaging and flow effects at these scales.

The physical/chemical consequences of fibrosis on liver performance are numerous. These include flow redistributions, changes in metabolic zonation and liver drug metabolizing enzymes, and drug propagation and effectiveness, (Dutta-Moscato et al, 2014; Ghallab A et al, 2019); Dhar et al, 2020). More particularly, Schenk et al (2017) address how resulting changes in enzyme zonation at the lobule level (“functional damage”) can impact reduced drug clearance capacity observed via blood plasma concentration profiles over time (using whole body physiologically-based pharmacokinetic PBPK modelling).

With the modern application of nanoparticle drug technology, the impact of fibrosis is even more consequential. Flow distribution of nanoparticles through a lobule is different from NP distribution into spheroids because of the presence of sinusoidal flow paths in the former. Here we have focussed on a specific paclitaxel-containing nanoparticle and highlighted the difference in behaviour of this NP with molecularly dissolved paclitaxel.

These simulations can be easily extended to other NP types and sizes by altering the NP diffusion and NP uptake and reaction parameters. Sofias et al (2017) gives a detailed comparison of paclitaxel-containing NPs.

The impact of collagen on nanoparticle distribution in tissue has been investigated experimentally by Diop-Frimpong et al (2011) and Sykes et al (2016). The first study indicated that the injection of a small-molecule collagen inhibitor (Losartan) improved the distribution and effectiveness of the Doxil liposomal nanoparticle. The second indicated that gold NP accumulation and penetration is altered by the tumour pathophysiology. Both NPs are of approximately 100 nm size. Simulations of these further effects employing our methods could be instructive.

Finally, it is emphasized that these simulations have assumed that the non-fibrotic lobule flow rates are maintained even under extensive fibrosis, and the net effect of fibrosis is an increase in lobule pressure gradients. This is often approximately true, but others have noted that fibrosis can also lead to somewhat reduced flow (and a smaller increase in lobule pressure gradients). In principle, the human body has other mechanisms to approximately maintain flow e.g. the use of vasoactive agents or a readjustment of hepatic artery to portal vein flow ratios. These effects are not within the scope of our individual lobule models, so we have chosen to maintain a constant flow boundary condition in our simulations. A flow reduction of e.g. 80% would result in a longer time production profile and less changes in lobule zonation pattern. Our earlier paper (Rezania et al, 2024) has analyzed this briefly.

Further resolution of the two somewhat competing consequences of fibrosis (flow rate reductions versus metabolic zonation changes at a fixed flow rate) can only be properly investigated using full multiscale modelling, transitioning from full body Physiologically Based Pharmacokinetic (PBPK) approaches through full liver vasculature to Capillary Surrogate Model (CSM) levels down to an individual lobule scale model such as ours. Here, an approach similar to that of Malka-Markovitz (2024) could be considered, based on a selected PBPK comparison of nanoparticle versus molecular dissolved drug delivery behaviour (Gabizon et al, 1994). The role of fibrosis on both these behaviours remains to be explored.

## References

Rezania, V, Coombe, D, Tuszynski, J, “A physiologically-based flow network model for hepatic drug elimination III: 2D/3D DLA lobule models”, Theor. Biol. Med. Model., v13, article 9 (2016);

Coombe, D, Wallace, C, Rezania, V, Tuszynski, J, “Computational analysis of upscaled fibrotic liver multi-lobule flows and metabolism”, MDPI Processes, v12, 1789, DOI.org10.3390 (2024);

Hulmes, D, “Quasi-hexagonal molecular packing in collagen fibrils”, Nature, v282, p878–880, (1979);

Hulmes, D, Wess, T, Prockop, D, Fratzl, P, “Radial packing, order, and disorder in collagen fibrils”, Biophys. J., v68, p1661–1670, (1995);

Orgel, D, Irving, T, Miller, A, Wess, T, “Microfibrillar structure of type I collagen in situ”, PNAS. V103, p9001–9005, (2006);

Silver, F, Freeman, J, Seehra, G, “Collagen self-assembly and the development of tendon mechanical properties”, Nature. v282, p878–880, (2003);

Buehler, M, “Atomistic and continuum modeling of mechanical properties of collagen: elasticity, fracture and self-assembly”, J., Mater. Res. v21, p1947–1961, (2006a);

Buehler, M, “Nature designs tough collagen: explaining the nanostructure of collagen fibrils”, PNAS., v103, p12285–12290, (2006b);

Vesentini, S, Redaelli, A, Gautieri, A, “Nanomechanics of collagen microfibrils”, Muscles, Ligaments and Tendons J., v3, p23–34, (2013);

Parkinson, J, Kadler, K, Brass, A, “Self-assembly of rodlike particles in two dimensions: A simple model for collagen fibrillogenesis”, Phys. Rev. E, v50, p2963–2950, (1994a);

Parkinson, J, Kadler, K, Brass, A, “Simple Physical Model of collagen fibrogenesis based on diffusion limited aggregation”, J. Mol. Biol., v247, p823–831, (1994b);

Parkinson, J, Brass, A, Canova, G, Brecht, Y, “Simple Physical Model of collagen fibrogenesis based on diffusion limited aggregation”, J. Biomech., v30, p549–554, (1997);

Garcia-Ruiz J, Otalora, F, “Diffusion limited aggregation. The role of surface diffusion”, Physica A, v178, p415–420, (1991);

Rothenbuhler, J, Huang, J, DiDonna, B, Levine, A, Mason, T, “Mesoscale structure of diffusion-limited aggregates of colloidal rods and disks”, Soft Matter, v5, p3639–3645, (2009);

Nicolas-Carlock, J, Carrillo-Estrada, J, Dossetti, V, “Fractality a la carte: a general particle aggregation model”, Sci. Reports, v6, 19505, DOI:10.1038, (2016);

Pederson, J, Boschetti, F, Swartz, M, “Effects of extracellular fiber architecture on cell membrane shear stress in a 3D fibrous matrix”, J. Biomech., v40, p1484–1492, (2007);

Stein, A, Vader, D, Jawerth, A, Weitz, D, Sander, L, “An algorithm for extracting the network geometry of three-dimensional collagen gels”, J. Microscop., v232 Pt 3, p463–475, (2008);

Lang, N, Munster, S, Metzner, C, Krauss, P, Schurmann, S, Lang, J, Aifantis, K, Friedrich, O, Fabry, B, “Estimating the 3D pore size distribution of biopolymer networks from directionally biased data”, Biophys. J., v105, p1967–1975, (2013);

Jackson, G, James, D, “The permeability of fibrous porous media”, Can. J. Chem. Eng., v64, p362–374, (1986);

Zhu, Z, Wang, Q, Wu Q, “On the examination of the Darcy permeability of soft porous media; new correlations”, Chem. Eng. Sc., v173, p525–536, (2017);

Bear, J, “Dynamics of fluid in porous media”, Elsevier, New York (1972);

Costa, A, “Permeability-porosity relationship: A reexamination of the Kozeny-Carmen equation based on a fractal pore-space geometry assumption”, Geophys. Res. Lett., v33, article L02318, (2006);

Ogston, A, Preston, Wells J, “On the transport of compact particles through solutions of chain polymers”, Proc. Roy. Soc. Lond. A, v333, p297–316, (1973);

Johnson, E, Berk, D, Jain, R, Deen, W, “Hindered diffusion of spherical macromolecules through dilute fibrous media”, Phys. Fluids., v8, p1720–1731, (1996);

Clague, D, Phillips, R, “A numerical calculation of the hydraulic permeability of three-dimensional disordered fibrous media”, Phys. Fluids, v9, p1562–1572, (1997);

Amsden, B, “Solute diffusion within hydrogels: mechanisms and models”, Macromolecules, v31, p8382–8395, (1998);

Phillips, R, “A hydrodynamic model for hindered diffusion of proteins and micelles in hydrogels”, Biophys. J., v79, p3350–3354, (2000);

Stylianopoulos, T, Diop-Frimpong, DB, Munn, L, Jain, R, “Diffusion anisotropy in collagen gels and tumors: the effect of fiber network orientation”, Biophys. J., v99, p3119–3128, (2010);

Gao, Y, Li, M, Chen, B, Sheng, Z, Guo, P, Wientjes, G, Au, J, “Predictive Models of Diffusive Nanoparticle Transport in 3-Dimensional Tumor Cell Spheroids”, AAPS J., v15, DOI: 10.1208/s12248-013-9478-2 (2013);

Gole, L, Liu, F, Ong, K, Li, L, Han, H, Young, D, Marini, G, Wee, A, Zhao, J, Rao, H, Yu, W, Wei, L, “Quantitative image-based collagen structural features predict the reversibility of hepatitis C virus-induced liver fibrosis post antiviral therapies”, Sc. Reports, v13, 6384, DOI:10.1038, (2023);

Keitzmann, T, “Metabolic zonation of the liver: The oxygen gradient revisited.”, Redox. Biol. v11, 622–630, (2017)

Ghallab, A, Myllys, M, Holland, C, Zaza, A, Murad, W, Hassan, R, Ahmed, Y, Abbas, T, Abdelrahim, E, Schneider, K, Matz-Soja, M, Reinders, J, Gebhardt, R, Berres, M, Hatting, M, Drasdo, D, Saez-Rodriguez, J, Trautwein, C, “Influence of Liver Fibrosis on Lobular Zonation”, Cells., v8, 1556, DOI:10.3390, (2019);

Kuh, H, Jang, S, Wientjes, G, Au, J, “Predictive Models of Diffusive Nanoparticle Transport in 3-Dimensional Tumor Cell Spheroids”, J. Pharmacol. Expt. Therap., v293, 761–770, (2000);

Vaclavikova, R, Soucek, P, Svobodova, L, Simek, P, Guengerich, F, Gut, I, “Different in vitro metabolism of paclitaxel and docetaxel in humans, rats, pigs, and minipigs”, Drug Metabolism and Disposition, v32(6), p666, (2004).

Nunes-Correia, I, Ramalho-Santos, J, Nir, S, Pedroso de Lima, M, “Interactions of Influenza Virus with Cultured Cells: Detailed Kinetic Modeling of Binding and Endocytosis”, Biochemistry., v38, 1095–1101, (1999);

Dutta-Moscato, J, Solovyev, A, Mi, Q, Nishikawa, T, Soto-Gutierrez, A, Fox, I, Vodovotz, Y, “A multiscale agent-based in silico model of liver fibrosis progression”, Front. Bioeng. Biotech., v2, article 18, (2014);

Dhar, D, Baglieri, J, Kisseleva, T, Brenner, D, “Mechanisms of liver fibrosis and its role in liver cancer”, Expt. Biol. Med., v245, p96–108, (2020);

Schenk, A, Ghallab, J, Hofmann, U, “Physiologically-based modelling in mice suggests an aggravated loss of clearance capacity after toxic liver damage”, Sci. Reports, v7, 6224, DOI:10.1038/s41598-017-0457/4-z (2017);

Sofias, A, Dunne, M, Storm, G, Allen, C, “The battle of nano paclitaxel”, Adv. Drug Delivery Rev., v122, p20–30, (2017);

Diop-Frimpong, B, Chauhan, V, Krane, S, Boucher, Y, Jain, R, “Losartan inhibits collagen I synthesis and improves the distribution and efficacy of nanotherapeutics”, PNAS., v108, p2909–2914, (2011);

Sykes, E, Dai, Q, Sarsons, C, Chen, J, Rocheleau, J, Hwang, D, Zheng, G, Cramb, D, Rinker, K, Chan, W, “Tailoring nanoparticle designs to target cancer based on tumor pathophysiology”, PNAS., v103, p12285–12290, (2016);

Malka-Markovitz, A, Dit Pinto, S, Cherkaoui, M, Levin, S, Anandasabapathy, S, Sood, G, Dhingra, S, Yujia, G, Vierling, J, Gallo, N, “Multiscale modeling of drug-induced liver injury from organ to lobule”, NPJ Digital Medicine., v8, #383, (2025);

Gabizon, A, Catane, R, Uziely, B, Kaufman, B, Safra, T, Cohen, R, Martin, F, Huang, A, Prolonged circulation time and enhanced accumulation in malignant exudates of doxorubicin encapsulated in polyethylene-gycol coated liposomes”, Cancer Res., v54, p987–993, (1994)

